# Importin α inhibitors act against the differentiated stages of apicomplexan parasites *Plasmodium falciparum* and *Toxoplasma gondii*

**DOI:** 10.1101/2024.07.10.602875

**Authors:** Manasi Bhambid, Sujata Walunj, Anupama C. A., Shilpi Jain, Diksha Mehta, Kylie Wagstaff, Ashutosh Panda, David A. Jans, Asif Mohmmed, Swati Patankar

## Abstract

Protozoan parasites of the phylum Apicomplexa, including *Plasmodium falciparum* and *Toxoplasma gondii*, cause widespread disease in humans. New drugs and protein targets are required for the treatment of these diseases, particularly therapies targeting multiple stages of the parasite life cycles. Nuclear import, carried out by the transporter’s importin (IMP) α and β subunits, is a valid target for the discovery of lead compounds against these protozoan parasites: small molecules were identified that inhibit interactions between IMPα and nuclear localisation signals *in vitro* and also inhibit the growth of the rapidly-dividing stages of *P. falciparum* and *T. gondii* (asexual stages and tachyzoites) in culture. In this report, we add another small molecule (Bay 11-7082) to the panel of inhibitors of IMPα and test the ability of these inhibitors to first, inhibit nuclear transport in the rapidly dividing stages and, next, the maturation of differentiated stages of both parasites. We show that GW5074 and CAPE inhibit nuclear transport in the *P. falciparum* blood stages, while Bay 11-7085 inhibits nuclear transport in *T. gondii* tachyzoites. Interestingly, CAPE strongly inhibits gametocyte maturation, the sexual stages of *P. falciparum*, and Bay 11-7085 weakly inhibits bradyzoite differentiation, the latent stages of *T. gondii*. As differentiation of both these stages is dependent on activation of gene expression, triggered by the nuclear translocation of transcription factors, our work provides a “proof of concept” that targeting nuclear import is a viable strategy for the development of therapeutics against multiple stages of apicomplexan parasites, some of them recalcitrant to existing drugs.

## Introduction

Apicomplexans are single-celled eukaryotes that cause disease ^1^. *Plasmodium* species cause malaria and of these, *P. falciparum* can give rise to fatal complications. Malaria burden has reduced worldwide, mainly due to artemisinin combination therapies (ACTs) ^2^. ACTs act on the asexual blood stages of the *Plasmodium* life cycle and some are effective against gametocytes, the sexual stages that drive transmission of malaria when taken up by mosquitoes ^2^.

Toxoplasmosis, caused by *Toxoplasma gondii*, affects one-third of the human population ^3^. *T. gondii* therapy combines antifolates, pyrimethamine and sulfadiazine ^4^. This therapeutic approach is effective against tachyzoite stages of the *Toxoplasma* life cycle but not against latent stages known as bradyzoites that form tissue cysts ^5^. There is an urgent need for new medications with novel mechanisms of action that can also target the differentiated stages of *P. falciparum* and *T. gondii*.

A handful of novel compounds act on the gametocytes of *P. falciparum* or the bradyzoites of *T. gondii* ^6–9^. The targets of these compounds include translation initiation, protein phosphorylation and other biological processes that play roles in differentiation. Another process that underlies differentiation is the activation of transcription factors, resulting in a cascade of gene expression. Indeed, in *P. falciparum*, PfAP2-G, an AP2 domain transcription factor, drives the progression of sexual stage parasites ^10,11^. Similarly, in *T. gondii*, the Bradyzoite-Formation Deficient 1 (BFD1) transcription factor triggers the differentiation of bradyzoites ^12^. For these transcription factors to carry out their functions, they must enter the nucleus.

Transport into the nucleus is mediated by the importin (IMP) superfamily of α and β proteins ^13^. In classical nuclear import, IMPα recruits IMPβ through its IMPβ-binding domain (IBB), and this complex binds the nuclear localisation signal (NLS) of cargo proteins to transport them into the nucleus ^14^. Dysregulation of nucleocytoplasmic transport can impact a range of cellular processes, including cell proliferation, growth and differentiation ^15,16^.

Blocking nuclear transport holds enormous potential for therapeutic intervention against pathogens. Compounds have been identified that block IMPα/β-dependent nuclear import of viral proteins central to infection in HIV, Zika virus, West Nile virus and Dengue virus ^17–19^. Similarly, general and specific inhibitors of IMPα from *P. falciparum* and *T. gondii* (PfIMPα, TgIMPα) reduce the growth of *P. falciparum* asexual stages and *T. gondii* tachyzoites ^20,21^. In this report, we add another compound (Bay 11-7082) to the panel of compounds and assess their ability to inhibit gametocytes and bradyzoites.

First, we show that Bay 11-7082, a chemically modified Bay 11-7085 analogue, binds directly to both PfIMPα and TgIMPα proteins, inhibiting their binding to NLSs *in vitro*. Next, we test the entire panel of small molecules for their ability to block nuclear transport in the asexual stages of *P. falciparum* and tachyzoites of *T. gondii*. GW5074 and CAPE inhibit nuclear transport in *P. falciparum*, while Bay 11-7085 inhibits nuclear transport in *T. gondii*. Consistent with their ability to block nuclear transport, CAPE inhibits gametocyte maturation, while Bay 11-7085 shows a small but statistically significant inhibition of tachyzoite to bradyzoite differentiation. This report provides “proof of concept” that targeting nuclear transport is a viable strategy for therapeutics against differentiated stages of *P. falciparum* and *T. gondii* and adds to the small but significant list of lead compounds that target these stages.

## Materials and Methods

### Protein expression and purification

PfIMPα, TgIMPα, MmIMPα, and MmΔIBBIMPα were expressed as GST fusion proteins and purified as described previously ^22,20,21^. His-tagged SV40 T-ag-NLS-GFP, TGS1-NLS-GFP, PfIMPα and TgIMPα proteins were purified using Ni^2+^-affinity chromatography as described previously ^23–25^.

### Compounds

Ivermectin, auranofin, GW5074, CAPE, Bay 11-7082 and Bay 11-7085 were purchased from Sigma-Aldrich, and dihydroartemisinin (DHA) was a kind gift from IPCA Laboratories, Mumbai. Stock solutions (10 mM) were prepared in 100% DMSO.

### AlphaScreen assay

The AlphaScreen assay was performed as described previously ^20,21^. The AlphaScreen signal was quantified on a Perkin Elmer plate reader, and titration curves (sigmoidal fit) were plotted using GraphPad Prism 9.0.2 (San Diego, California, USA).

### Intrinsic tryptophan fluorescence assays

Experiments were performed using a JASCO Fluorescence Spectrophotometer and 0.5 mL quartz cuvettes, with a slit width of 5 nm at a constant temperature of 25°C. Emission spectra were collected between 310 and 400 nm after excitation at 295 nm ^26^. The blank correction was performed using 1X PBS. Bay 11-7082 and Bay 11-7085 (5-80 μM) were incubated with 1 μM IMPα proteins for 5 minutes at room temperature, with DMSO being the control. Data from three experiments were analysed using GraphPad Prism 9.0.2 (San Diego, California, USA).

### Cloning of PTEF-NLS-GFP and GFP-SV40 T-ag-NLS-GFP

The codon-optimized NLS of the *Plasmodium* Translation-Enhancing Factor (PTEF; PlasmoDB ID: PF3D7_0202400) was cloned into pSSPF2 ^27^, using annealed primers with BglII and AvrII sites (bold). Primers are shown below, having the NLS sequence (underlined) and a preferred Kozak sequence (italicised) ^28^.

PTEF-NLS primer 1: 5ʹATA**AGATCT***AAAAAATGG*GA<u>AAAAAGAACAGAGATAAGAAACATTCTAAAAAAAGGAAAACA</u> <u>AAACAAAATTATAAA</u>**CCTAGG**AGT3ʹ

PTEF-NLS primer 2: 5ʹACT**CCTAGG**<u>TTTATAATTTTGTTTTGTTTTCCTTTTTTTAGAATGTTTCTTATCTCTGTTCTTTT</u>T TCCCATTTTTT**AGATCT**TAT3ʹ

For GFP-SV40 T-ag-NLS-GFP, a second Green Fluorescent Protein (GFP) gene was cloned into pCTG-eGFP using the NheI and AvrII sites (bold). The forward primer harboured the preferred *Toxoplasma* Kozak sequence (italicised) and the start codon ^29^. Next, the SV40 T-ag-NLS (underlined) was cloned in-frame between the two GFP genes.

GFP-GFP forward:

5ʹAAG**GCTAGC***GACAAA*ATGGTGAGCAA3ʹ

GFP-GFP reverse:

5ʹAAA**CCTAGG**CTTGTACAGCTCGTCCAT3ʹ

GFP-SV40 T-ag-NLS-GFP forward: 5ʹ<u>CCCAAGAAAAAACGCAAGGTG</u>GTGAGCAAGGGCGAGGAGCTGTTC3ʹ

GFP-SV40 T-ag-NLS-GFP reverse: 5ʹCTTGTACAGCTCGTCCATGCCGAGAGTGATC3ʹ

All primers were obtained from Sigma-Aldrich, and inserts were confirmed by sequencing to be free of mutations.

### *P. falciparum* and *T. gondii* culture and transfections

*P. falciparum* (3D7 strain) was cultured using standard procedure ^30^. For stable transfections, parasites were synchronised using 5% D-sorbitol to obtain early-stage rings at 5% parasitemia. For electroporation using the Bio-Rad GenePulser Xcell system, 200 μl of parasites were mixed with 100 μg of purified PTEF-NLS-GFP plasmid and 1X cytomix in a 2 mm cuvette (settings: 0.310 kV, 950 μF, ∞ Ω, 2 mm cuvette) ^31^. Transgenic parasites were selected with 2.5 μg/ml Blasticidin ^32^ over 3-4 weeks. The stable line was synchronised (4-5% parasitemia, late rings to early trophozoites) before treatment with small molecules.

*T. gondii* RH and ME49ΔKu80 strains were maintained in Human Foreskin Fibroblast (HFF) cells (ATCC) using standard culture conditions ^33^. For transient transfections, 2 x 10^6^ RH parasites and 50 μg of GFP-SV40 T-ag-NLS-GFP plasmid with a final concentration of 1X cytomix were electroporated (1500 V, 25 μF, 50 Ω, 2 mm cuvette) ^33^. After 4-5 hours, parasites were treated with compounds ^20,21^.

### Drug susceptibility assays for parasites and cytotoxicity assay for HFF cells

*P. falciparum* cultures (3% hematocrit, 5% CO_2_) were synchronised (0.25% parasitemia, rings). IC_50_ was determined as previously reported using the HRP2-sandwich ELISA assay ^20,21^. The *T. gondii* RH strain constitutively expressing a luciferase reporter was used for the drug susceptibility assays as previously described ^20^. Luminescence was measured using a Varioskan LUX multimode microplate reader (Thermo Fisher Scientific). Cytotoxicity of the small molecules for the host HFF cells (100% confluent monolayer) was assayed using the MTT assay as previously described ^34^.

### Immunofluorescence assays and image analysis

After 10 hours of drug treatment, *P. falciparum* parasites (trophozoites) were fixed in the dark with 4% paraformaldehyde (Merck) and stained with DAPI (Sigma-Aldrich) (5 μg/ml). Images were acquired (50-60 per sample) by Nikon A1 confocal laser scanning microscope with a 100X objective. A region of interest was drawn around the DAPI signal in the entire stack, and colocalisation with GFP was analysed using Pearson’s correlation coefficient (PCC) with the JaCoP plugin in ImageJ (version 2.0.0) ^35^. Dot plots of PCC values were plotted in GraphPad Prism (version 9.5.1) (San Diego, California, USA), and statistical analysis was done using the Mann-Whitney test. The three-dimensional images were constructed using IMARIS software (version 8.3.1) (RRID: SCR_007370).

For *T. gondii* tachyzoites, after 24 hours of drug treatment, intracellular parasites were fixed with 4% formaldehyde (Merck) and 0.0075% glutaraldehyde (Sigma-Aldrich) in the dark. Parasites were permeabilised with 0.25% Triton X-100 (Sigma-Aldrich) and stained with DAPI (Sigma-Aldrich) (2 μg/ml). Parasite images (25-30 per sample) were acquired using a Zeiss LSM 780 confocal microscope and analysed for the ratio of nuclear signal to cytoplasmic signal intensity of GFP using the CellProfiler software (version 4.2.6) ^36^. Dot plots and statistical analysis were done as described earlier.

### Gametocyte induction

A clinical sample that spontaneously forms gametocytes was cultured in RPMI media (Invitrogen) complemented with 0.5% (w/v) Albumax II (Invitrogen) at 4% hematocrit and grown at 37°C in a gas mixture ^37^. Nutrient stress was started at high parasitemia (daily 80% fresh media added and no fresh RBCs) ^38,39^, and within 3-5 days, early gametocytes (Stage I/II) were observed. N-acetyl glucosamine (HiMedia) (50 mM) was added for 72 hours to block merozoite invasion ^39^. At 4% gametocytaemia, early to mid-stage gametocytes (Stage III/IV) were treated with compounds at their IC_50_ concentration (in the 3D7 strain) and 0.3% DMSO was used as the control.

After 48 and 72 hours of treatment, smears were made from the culture and stained with modified Giemsa stain (Sigma-Aldrich). The % (per total RBCs) of mid and mature gametocytes was counted ^38,39^. Bar graphs were plotted in GraphPad Prism (version 9.5.1) (San Diego, California, USA), and statistical analysis was done using the unpaired t-test.

### Bradyzoite Induction

Confluent HFF cells were infected with ME49ΔKu80 parasites (1 parasite: 5 hosts) in DMEM media (Invitrogen) with 10% (v/v) FBS (HiMedia), pH 7.4. The culture was incubated at 37°C and 5% CO_2_ for 4-5 hours to allow invasion. The media was changed to bradyzoite-inducing alkaline media (DMEM, 50 mM HEPES, 10% FBS, pH 8.2) ^40^. The parasites were incubated at 37°C, ambient CO_2_ to allow bradyzoite differentiation.

Compounds (IC_50_ concentrations from the RH strain) were added either with the alkaline media or when the mature cysts formed after 48 hours of alkaline media treatment. Treatment was continued for 24 hours, with a 0.1% DMSO control included.

### DBA Lectin Immunofluorescence and Image Analysis

After 24 hours of treatment, cells were fixed, permeabilised as described earlier, and stained with DAPI and Dolichos Biflorus Agglutinin-Lectin (DBA-Lectin) conjugated to FITC (Invitrogen) (16 μg/mL) for 1 hour. The slides were visualised with a Zeiss observer Z1 Spinning Disc microscope. Image analysis using Colocalisation Image Creator and Colocalisation Object counter plugins was done in ImageJ (version 2.0.0) ^41^. Cysts had nuclear (DAPI) signals colocalising with DBA-Lectin:FITC, while HFF cells showed only DAPI staining. The number of cysts in each field and the total DAPI signals in 15-20 fields were counted. Graphs were plotted in GraphPad Prism (Version 8.4.3) (San Diego, California, USA) after calculating the percentage of bradyzoite cysts per HFF in a field. Statistical analysis was done using the Mann-Whitney test.

## Results and Discussion

### Bay 11-7082 binds to PfIMPα and TgIMPα, inhibits interactions with NLSs and viability of *T. gondii* in culture

Previously, Bay 11-7085 showed stronger inhibition of the interaction between PfIMPα and TGS1-NLS than its inhibition of the interactions between TgIMPα or MmΔIBBIMPα with the SV40 T-ag NLS _20_. Further, thermal stability assays showed that Bay 11-7085 stabilised PfIMPα but not the other two IMPα proteins. These data suggested that Bay 11-7085 may bind to PfIMPα at sites distinct from those on TgIMPα and MmIMPα _20_.

Given these interesting results, we studied Bay 11-7082, a Bay 11-7085 analogue, having a single methyl group in its structure, whereas Bay 11-7085 has three ^42^ (Figure 1 a). In contrast to Bay 11-7085, Bay 11-7082 inhibited both PfIMPα:NLS and TgIMPα:NLS binding with IC_50_ values of 3.9 ±0.1 μM and 8.2 ± 1.3 μM, respectively (Figure 1 b, Table 1). Bay 11-7082 also inhibited MmΔIBBIMPα:NLS with IC_50_ values of 6.7 ± 2.1 μM (Figure 1 b, Table 1). Therefore, compared to Bay 11-7085, the single methyl group in the structure of Bay 11-7082 expands its specificity.

**Table 1.** Data from *in vitro* and *in vivo* assays for Bay 11-7082 and Bay 11-7085 compounds. T-ag-NLS: SV40 T-ag-NLS; HFF: Human foreskin fibroblast cells; Not measurable: curve fitting did not yield a measurable K_D_ value in the concentration range tested; No killing: parasite killing not observed till 100 μM. Cell viability assays were done on *P. falciparum* asexual blood stages and *T. gondii* tachyzoite stages.

| Bay 11-7082 |  |  | Bay 11-7085 |  |  |
| --- | --- | --- | --- | --- | --- |
| AlphaScreen |  |  |  |  |  |
| Proteins | IC<sub>50</sub> (μM) | Reference | Proteins | IC<sub>50</sub> (μM) | Reference |
| PfIMPα:<br>TGS1-NLS | 3.9 ± 0.1 | This report | PfIMPα:<br>TGS1-NLS | 1.9 ± 0.1 | 20 |
| TgIMPα:<br>T-ag-NLS | 8.2 ± 1.3 |  | TgIMPα:<br>T-ag-NLS | 30.4 ± 0.6 |  |
| MmΔIBBIMPα:<br>T-ag-NLS | 6.7 ± 2.1 |  | MmΔIBBIMPα:<br>T-ag-NLS | 23.4 ± 2.9 |  |
| Tryptophan fluorescence |  |  |  |  |  |
| Proteins | IC<sub>50</sub> (μM) | Reference | Proteins | IC<sub>50</sub> (μM) | Reference |
| PfIMPα | 21 ± 4.9 | This report | PfIMPα | 19.9 ± 3.7 | This report |
| TgIMPα | 17.5 ± 3.3 |  | TgIMPα | 27.7 ± 9.4 |  |
| MmIMPα | Not measurable |  | MmIMPα | Not measurable |  |
| Cell viability |  |  |  |  |  |
| Organism | IC<sub>50</sub> (μM) | Reference | Organism | IC<sub>50</sub> (μM) | Reference |
| *P. falciparum* | No killing | This report | *P. falciparum* | No killing | 20 |
| *T. gondii* | 1.06 ± 0.07 |  | *T. gondii* | 2.7 ± 0.7 |  |
| HFF | 12.4 ± 0.32 |  | HFF | 8.2 ± 0.7 |  |

**Figure 1.**
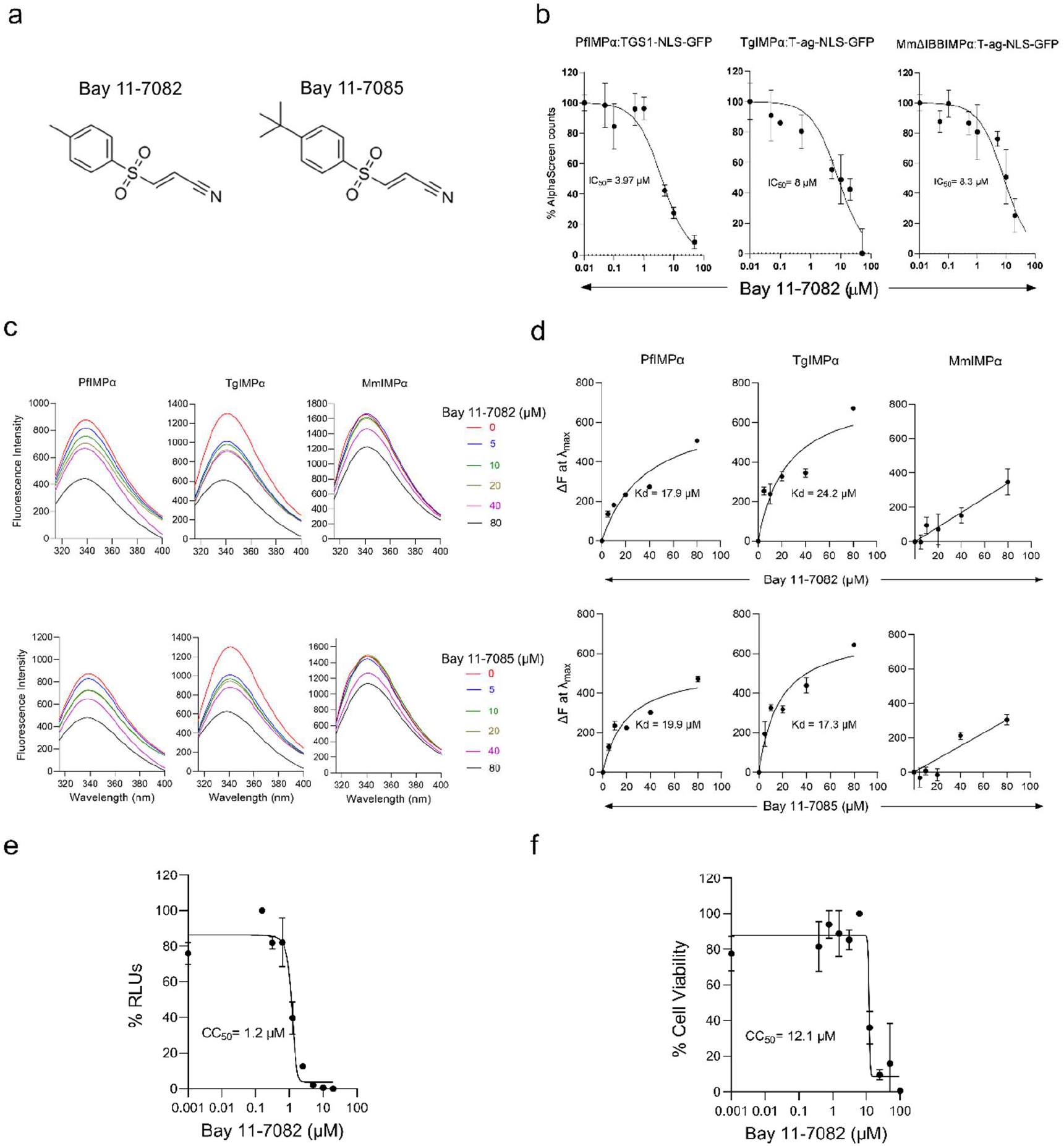
Effect of Bay 11-7082 on IMPα proteins and cells. **(a)** Structure of Bay 11-7082 and Bay 11-7085 from PubChem (https://pubchem.ncbi.nlm.nih.gov); **(b)** Bay 11-7082 inhibits interactions between apicomplexan IMPα and NLS as shown by AlphaScreen. AlphaScreen technology was used to determine the IC_50_ for inhibition by Bay 11-7082 of IMPα (5 nM) to NLS (30 nM). Data represent the mean ± SEM (n = 4) from a single experiment from a series of 3 independent experiments; **(c)** Intrinsic tryptophan fluorescence spectra of PfIMPα, TgIMPα and MmIMPα were collected in the presence or absence of Bay 11-7082 and Bay 11-7085 at indicated concentration ranges; **(d)** Compound concentrations vs changes in fluorescence intensity with Bay 11-7082 and Bay 11-7085 at a single fluorescence wavelength (*λ*max) were plotted using GraphPad prism. Data were plotted as mean ± SD from three independent experiments for each assessment; **(e)** Cell viability assay of *T. gondii* tachyzoites when treated with Bay 11-7082. RLUs represent the relative light units from a strain stably expressing firefly luciferase; **(f)** MTT assay of HFF cells.

Intrinsic tryptophan fluorescence was used to confirm these results. Tryptophan residues emit fluorescence, and fluorescence intensity is altered in the case of ligand binding ^26,43^. Interestingly, in importin α proteins, tryptophan residues are only present in the NLS-binding sites and altered intensity would indicate ligand binding in these sites.

Emission spectra of the IMPα proteins showed a fluorescence maximum at 340 nm (Figure 1 c). Both Bay compounds caused a concentration-dependent decrease in fluorescence intensity at 340 nm for PfIMPα and TgIMPα (Figure 1 c). Using these data, IC_50_ values were calculated. Bay 11-7082 and Bay 11-7085 had IC_50_ values for binding to PfIMPα of 21 ± 4.9 μM and 19.9 ± 3.7 μM, respectively and IC_50_ values for binding to TgIMPα of 17.5 ± 3.3 μM and 27.7 ± 9.4 μM, respectively (Figure 1 d, Table 1). Both the compounds did not show significant binding to full-length MmIMPα (Figures 1 c, d, Table 1). These results indicate that the compounds have a higher affinity for apicomplexan importin α proteins. As IMPα proteins have tryptophan residues only in their NLS-binding sites, the data also suggest that in the NLS-binding assays, Bay 11-7082 inhibits NLS interactions with PfIMPα and TgIMPα by direct binding to the proteins and inhibits NLS interactions with MmIMPα by binding to the NLS.

Parasite viability assays were carried out for Bay 11-7082 using *P. falciparum* asexual stages (3D7 strain) and *T. gondii* tachyzoites (RH strain). Bay 11-7082 did not inhibit *P. falciparum* growth (data not shown), similar to Bay 11-7085 ^20^, indicating cell-permeability issues. Bay 11-7082 inhibited *T. gondii* tachyzoites with an IC_50_ of 1.06 ± 0.07 μM (Figure 1 e, Table 1). Interestingly, when tested in an MTT assay using HFF cells, Bay 11-7082 showed a CC_50_ of 12.04 ± 0.32 μM (Figure 1 f, Table 1). This selectivity index (SI) of 11.4 is higher than the SI of 3.04 seen for Bay 11-7085 ^20^ and is consistent with the tryptophan fluorescence assays showing binding of Bay 11-7082 to TgIMPα and not to MmIMPα.

### GW5074 and CAPE inhibit nuclear localisation in *P. falciparum* asexual cultures

In order to draw a direct link between *in vitro* assays and growth inhibition, we tested the ability of the panel of compounds to inhibit nuclear localisation in *P. falciparum*. The PTEF-NLS was chosen as it was shown to carry GFP to the nucleus in a previous report ^44^. A stable *P. falciparum* line was generated that expressed the PTEF-NLS-GFP fusion protein in the nucleus and the cytosol (Figure 2 a, b) under control conditions (DMSO). Treatment with GW5074 and CAPE resulted in a statistically significant reduction in colocalisation of NLS-GFP with DAPI (Figure 2 a, b, c) compared to the control. Therefore, these compounds inhibit nuclear transport *in vivo* despite showing no structural similarity (Figure 2 d). As GW5074 and CAPE inhibit both PfIMPα:NLS binding *in vitro* and localisation of PTEF-NLS-GFP into the nucleus, they are authentic inhibitors of nuclear transport in *P. falciparum* asexual stages. Of these compounds, GW5074 was shown to block the nuclear localisation of a viral NLS in human cells at 20 μM ^45^, while the effects of CAPE on nuclear import have been reported here for the first time for a human pathogen.

**Figure 2.**
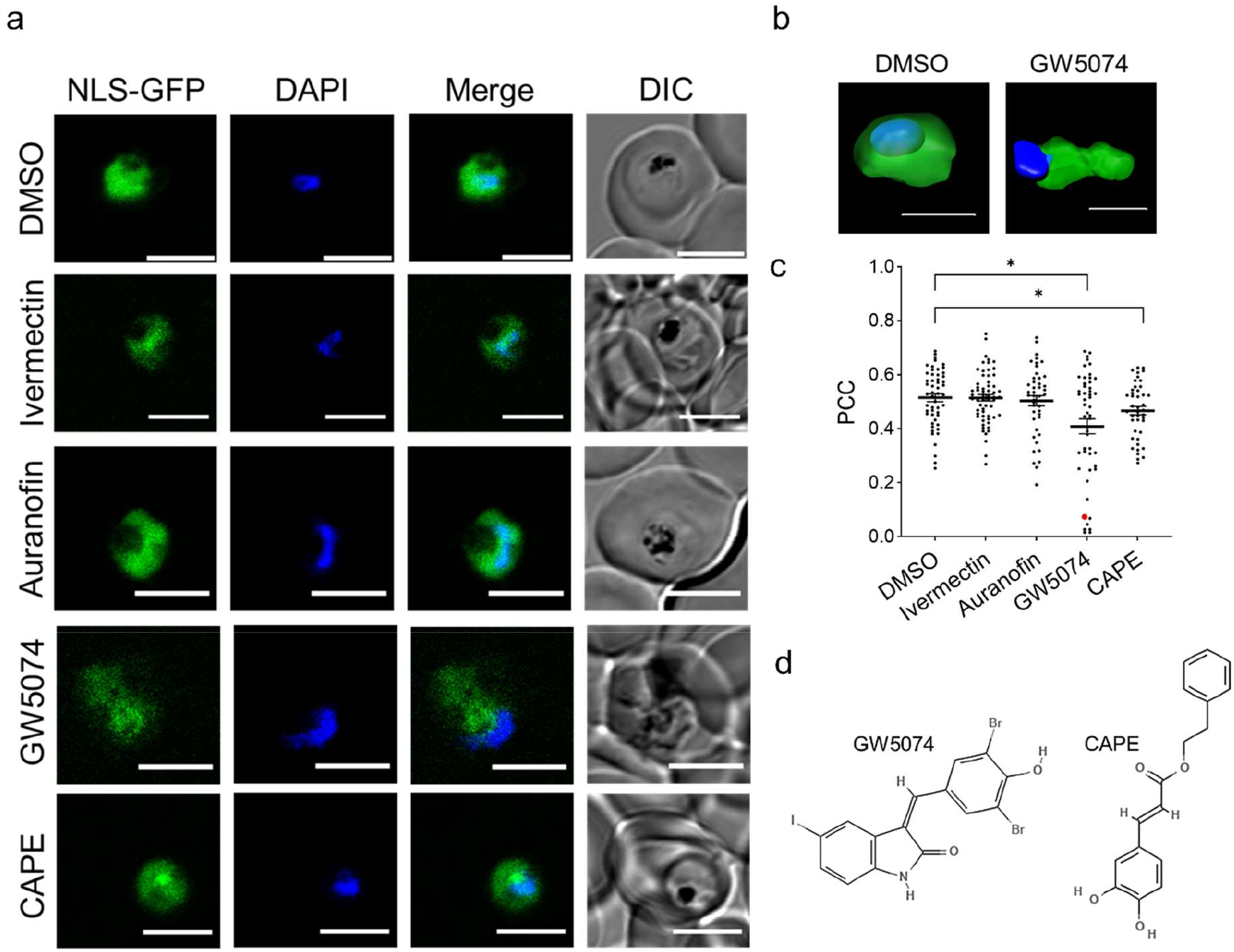
GW5074 and CAPE affect NLS-GFP nuclear localisation in *P. falciparum* blood stages. **(a)** PTEF-NLS-GFP localises to the nucleus and cytosol in the trophozoite stages. Parasites were treated for 10 hours at IC_50_ concentrations and imaged at 100X magnification using a laser scanning microscope. Scale bar: 5 microns; **(b)** 3D reconstruction of the parasites from panel (a) treated with GW5074 and DMSO control using Imaris software; **(c)** Colocalisation of PTEF-NLS-GFP with DAPI is plotted as a median with 95% CI, with each dot representing a parasite. * indicates a significant difference of p < 0.05. The red dot is the parasite in the 4^th^ row of the image panel; **(d)** Structure of GW5074 and CAPE from PubChem.

### CAPE inhibits the differentiation of mid-stage and mature gametocyte stages of *P. falciparum*

Differentiation and maturation of *P. falciparum* gametocytes is dependent on PfAP2-G ^11^. This protein appears to be a cargo for PfIMPα as the cNLS Mapper software (https://nls-mapper.iab.keio.ac.jp/cgi-bin/NLS_Mapper_form.cgi), trained to identify classical NLSs recognised by IMPα _46,47_, identifies a monopartite NLS starting at amino acid 471 (score of 6) and a bi-partite NLS starting at amino acid 446 (score of 6.6). Therefore, we tested whether the panel of compounds that inhibit PfIMPα (ivermectin, auranofin, GW5074 and CAPE) could also inhibit the differentiation of gametocytes. The anti-malarial drug dihydroartemisinin (DHA) was taken as a control.

A clinical strain of *P. falciparum* was induced to form gametocytes by nutrient stress (Figure 3 a), which resulted in cultures enriched in mid-stage (Stage III/IV) and late-stage (Stage V) gametocytes (Figure 3 b). Unhealthy gametocytes (Figure 3 c), ranging from 0.5 to 1% parasitemia, were seen in all the wells but not included in the data analysis. However, treatment with ivermectin resulted in a two-fold increase in unhealthy gametocytes (data not shown).

**Figure 3.**
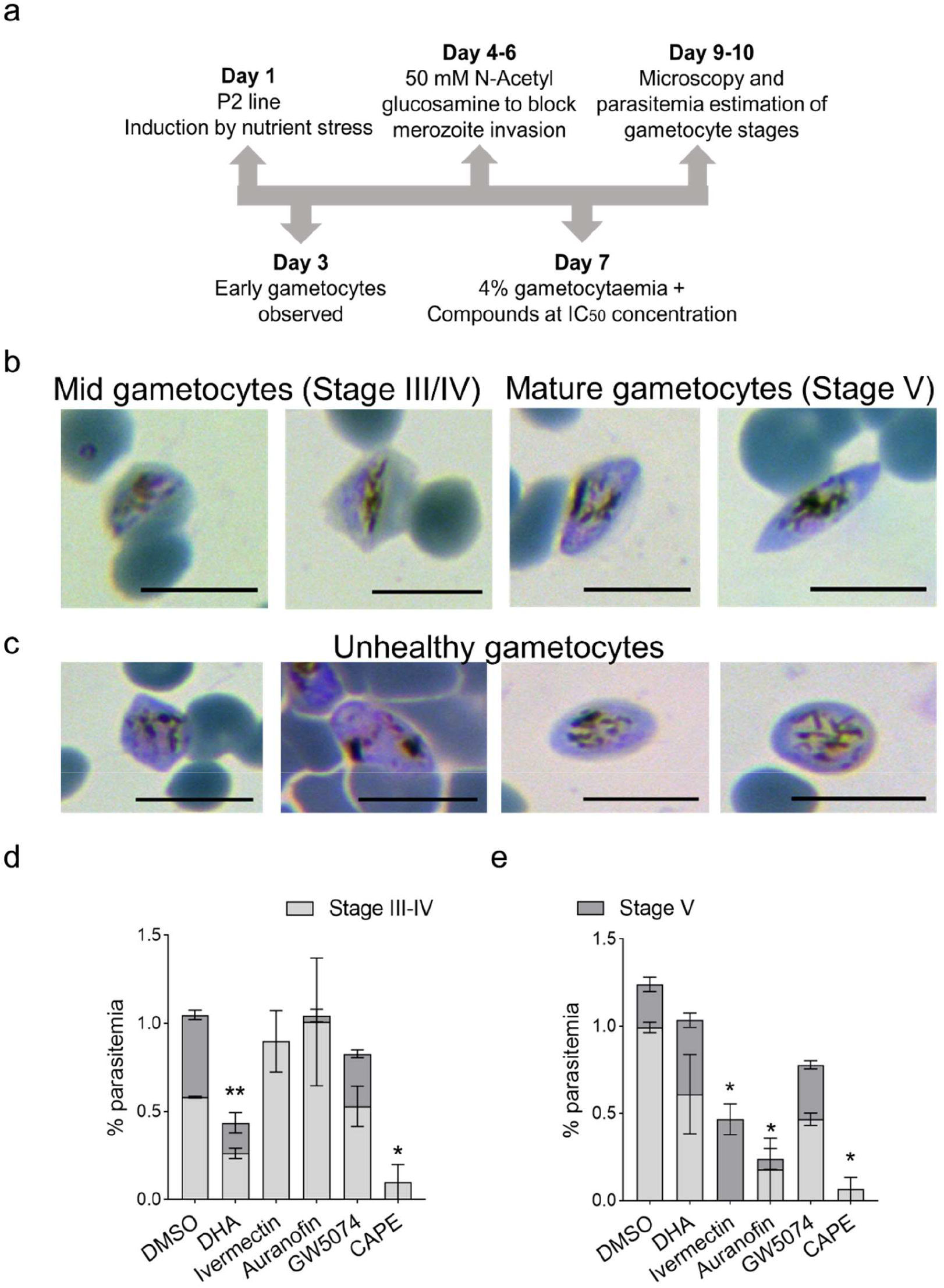
Gametocyte viability is severely affected by auranofin and CAPE. **(a)** Schematic for gametocyte induction and viability assay of *P. falciparum* clinical isolate Patient 2 (P2) when induced by nutrient stress ^39^; **(b)** Images of healthy Stage III/IV (8-10 days post-induction) and Stage V (11-13 days post-induction) seen by microscopy ^53^; **(c)** Unhealthy gametocytes were observed in culture and were not included in the estimation. Scale bar: 5 microns; % parasitemia of gametocytes treated for **(d)** 48 and **(e)** 72 hours at IC_50_ concentration of small molecules. % parasitemia is the number of parasites per 100 RBCs and plotted as mean ± SEM for two technical replicates. * indicate a significant difference of p < 0.05.

After 48 hours of treatment, compared to the DMSO control, DHA and CAPE showed a statistically significant reduction in the percentage of mid-stage and late-stage gametocytes (Figure 3 d). After 72 hours of treatment, DHA showed a similar percentage of mid-stage and late-stage gametocytes as the control (Figure 3 e), consistent with published literature that this anti-malarial drug does not affect late-stage gametocytes ^48^. Ivermectin and auranofin decreased Stage V gametocytes, suggesting they may affect sexual stage maturation by blocking pathways other than nuclear transport. Interestingly, the dramatic reduction in mature gametocytes continued with CAPE treatment after 72 hours (Figure 3 e). Therefore, of the two authentic inhibitors of PfIMPα identified in the previous section (Figure 2), CAPE is able to significantly inhibit gametocyte maturation in culture.

There are reports of CAPE blocking nuclear translocation of the NF-κB subunit at 20 μM, but not of other transcription factors ^49,50^, indicating specificity towards NLSs. Thus, CAPE (not GW5074) may have blocked the interactions of PfIMPα with gametocyte-specific transcription factors such as PfAP2-G. These data suggest that novel lead compounds targeting PfIMPα might be discovered if the experimental design for *in vitro* screening uses different NLS sequences. Further validation of CAPE’s cellular target(s) is needed, including experiments such as selecting drug-resistant parasites in culture, followed by genome sequencing. Other approaches could involve defining changes in the *P. falciparum* nuclear proteome after CAPE treatment.

### Bay 11-7085 inhibits nuclear transport in *T. gondii* tachyzoites

Similar to the experiments done in *P. falciparum*, we tested whether Bay 11-7082, Bay 11-7085 and auranofin can block nuclear transport in *T. gondii* tachyzoites. We encountered unexpected technical issues while expressing fusion proteins of different NLSs with GFP reporters in *T. gondii* (data not shown). After empirically testing several fusion proteins, we found that the SV40 T-ag-NLS, when placed between two GFP reporter proteins and expressed transiently, showed a GFP signal in both the nucleus and cytoplasm (Figure 4 a).

**Figure 4.**
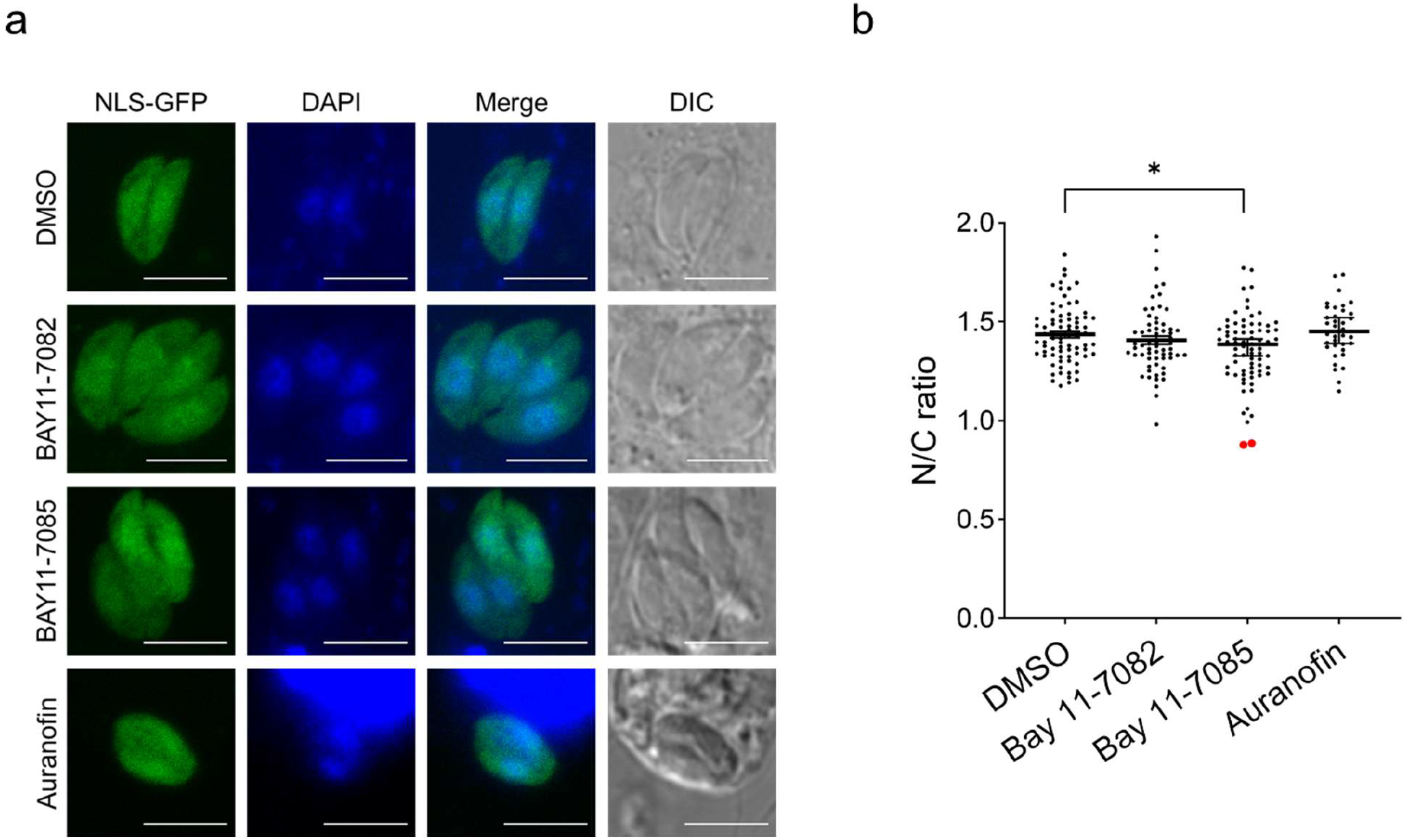
Bay 11-7085 affects the GFP-NLS-GFP localisation to the nucleus of *T. gondii* tachyzoites. **(a)** GFP-SV40 T-ag-NLS-GFP localises to the nucleus and cytosol in the tachyzoites. Parasites were treated for 24 hours at IC_50_ concentrations and imaged at 100X magnification using a laser scanning microscope. Scale bar: 5 microns; **(b)** Nuclear to cytoplasmic ratio of GFP-SV40 T-ag-NLS-GFP signal in the cell is plotted as a median with 95% CI, with each dot representing a parasite. * indicates a significant difference of p < 0.05. The red dots are the parasites in the 3^rd^ row of the image panel with the lowest N/C ratio.

IC_50_ concentrations of compounds resulted in the nuclear to cytoplasmic ratio of the GFP signal being significantly decreased with Bay 11-7085 (Figure 4 a, b). As Bay 11-7085 inhibits the interaction between TgIMPα and SV40 T-ag-NLS in the AlphaScreen assay ^20^ and inhibits nuclear localisation of this NLS *in vivo*, it is an authentic inhibitor of nuclear transport in *T. gondii* tachyzoites.

### Bay 11-7085 affects the number of *T. gondii* bradyzoite cysts in culture

Bradyzoites are latent stages of the *T. gondii* life cycle, dependent, among other factors, upon the transcriptional activation of genes by BFD1 ^12^. cNLS mapper identified a monopartite NLS in BFD1 at amino acid 1244 (score of 10), suggesting its nuclear translocation could be TgIMPα-dependent.

By alkaline stress, ME49ΔKu80 tachyzoites were induced to differentiate into bradyzoites with cyst walls that bind to fluorescently labelled DBA-lectin (Figure 5 a, b). Cultures were treated with IC_50_ concentrations of the anti-*Toxoplasma* drug pyrimethamine, Bay 11-7082, Bay 11-7085 or auranofin, with the treatment regimen initiated either at the start of bradyzoite induction or after mature bradyzoites were observed (summarised in Figure 5 a). Before starting these experiments, the IC_50_ values of the compounds in alkaline media were assayed on the RH-Fluc strain as described previously ^20^, and no significant differences from the values obtained at pH 7.4 were observed (data not shown).

**Figure 5.**
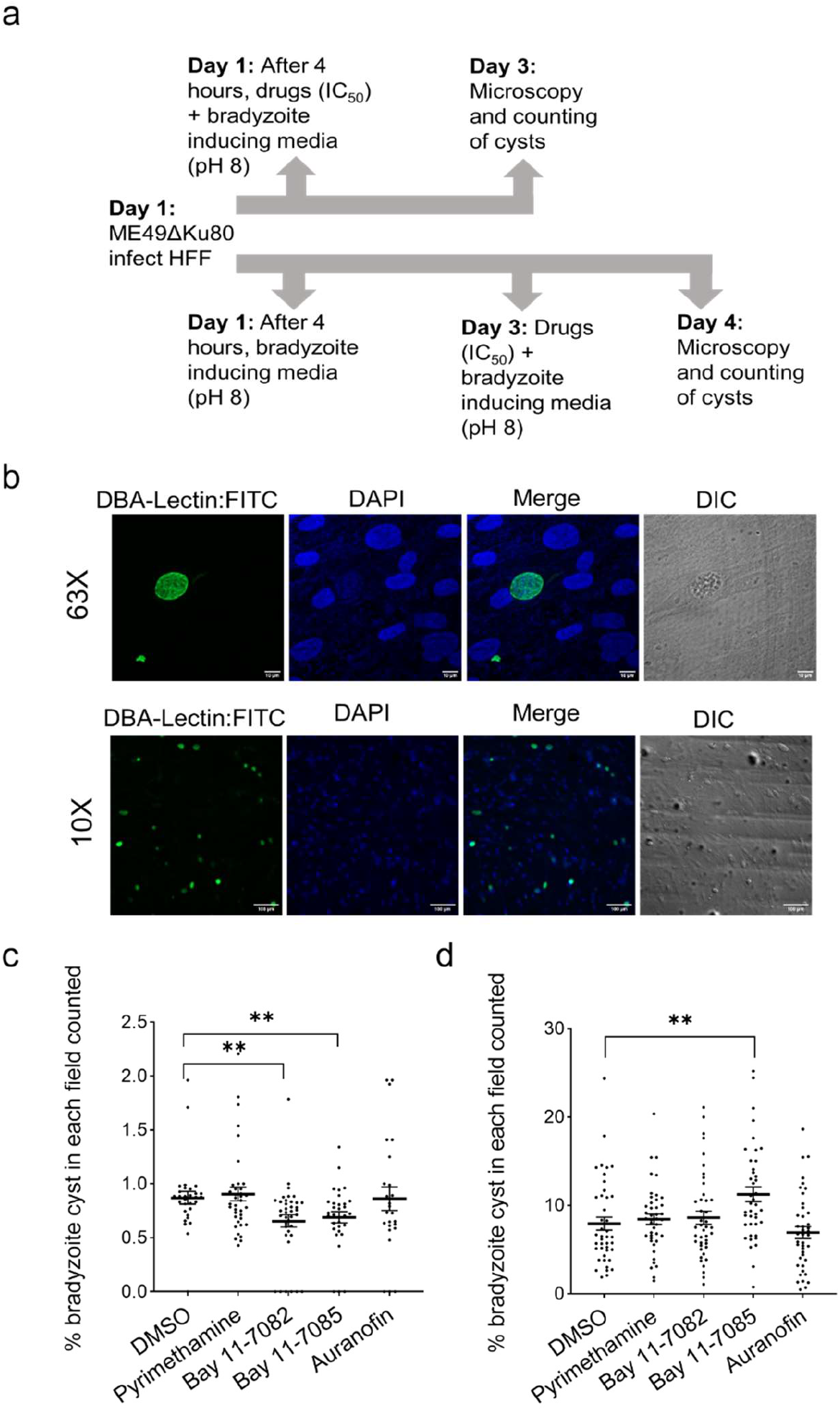
Bay 11-7085 affects tachyzoite to bradyzoite stage conversion. **(a)** Schematic of drug assay. Compounds were added at the same time as alkaline stress or after allowing bradyzoite conversion for 48 hours. The alkaline pH was maintained till the end of the experiments and compounds were added to the fresh media; **(b)** Bradyzoites (DMSO control) visualised using confocal microscopy after DBA-lectin:FITC and DAPI staining with 63X and 10X magnification; **(c)** Effect of drugs on bradyzoite differentiation when added at the tachyzoite stages; **(d)** Effect of drugs on differentiated bradyzoite cysts. For (c) and (d), data were obtained at 24 hours post-treatment and are plotted as a percentage of bradyzoite cysts counted per field (number of bradyzoites [FITC colocalising with DAPI]/total number of cells [DAPI] x100). * indicates a significant difference of p < 0.05 compared to vehicle control. Note that only fields having > 100 HFF cells were included in the analysis.

Treatment of tachyzoites with Bay 11-7082 and Bay 11-7085 showed a statistically significant decrease in bradyzoite numbers at 24 hours post-treatment (Figure 5 c). These early results hold promise for testing other analogues with different chemical modifications of the scaffold ^42,51^. In order to block bradyzoite formation, small molecules must inhibit interactions between the BFD1-NLS and TgIMPα. Therefore, performing HTS of chemical libraries using the BFD1-NLS rather than the SV40 T-ag-NLS might identify novel compounds that block bradyzoite differentiation.

When mature bradyzoites were treated with the compounds, Bay 11-7085 resulted in an increase in bradyzoite numbers (Figure 5 d), appearing to induce a stress response that supersedes its ability to block nuclear transport. Importantly, small molecules must penetrate the cyst wall of bradyzoites, and novel methods for delivery of Bay 11-7085 may need to be developed ^52^. Further validation of TgIMPα as the target of Bay 11-7085 *in vivo* will confirm these preliminary findings.

In conclusion, the pipeline for combating malaria and toxoplasmosis needs new lead compounds with novel mechanisms of action, especially those that target different stages of the life cycles of these widespread human pathogens. This report identifies such lead compounds, providing a “proof of principle” for the first time that targeting nuclear import can block many stages of these apicomplexan parasites.

### Ethics approval

Blood for *P. falciparum* culture was obtained from volunteers, and the gametocyte line (P2) of *P. falciparum* was obtained from a patient after approval from the Institute Ethics Committee, AIIMS, India (IEST/T-438/30.11.2012).

## Acknowledgements

We acknowledge Anjali Arya for doing the *T. gondii* drug assays at pH 8.2. We thank the Indian Institute of Technology Bombay (IIT Bombay) for their Confocal Laser Scanning Microscope and Spinning Disc Confocal Microscope, and the ICGEB, New Delhi, for their Confocal Laser Scanning Microscope. We thank Sanjeeva Srivastava and Prakriti Tayalia, IIT Bombay, for using plate readers for the parasitic drug assays and the IIT Bombay Hospital staff and volunteers for their help in blood collection.

## Author contributions

SW, DM, SP, KW and DAJ conceived and designed the experiments for Figure 1. MB, SJ and AM conceived and designed the experiments for Figures 2 and 3. AP generated the clinical strain for Figure 3. MB and SP conceived and designed the experiments for Figure 4. ACA and SP conceived and designed the experiments for Figure 5. MB, SW, SJ, ACA and DM performed the experiments. MB, SW, SJ, ACA and DM prepared the figures. MB and SP prepared the manuscript. AM and DAJ gave feedback on the drafts of the manuscript and all the authors reviewed the final manuscript.

## Transparency declaration

The authors declare they have no competing interests.

## References

1. Adl, S. M. et al. Diversity, Nomenclature, and Taxonomy of Protists. Systematic Biology 56, 684–689 (2007).

2. Greenwood, B. M. et al. Malaria: progress, perils, and prospects for eradication. J. Clin. Invest. 118, 1266–1276 (2008).

3. Tenter, A. M., Heckeroth, A. R. & Weiss, L. M. Toxoplasma gondii: from animals to humans. International Journal for Parasitology 30, 1217–1258 (2000).

4. Dunay, I. R., Gajurel, K., Dhakal, R., Liesenfeld, O. & Montoya, J. G. Treatment of Toxoplasmosis: Historical Perspective, Animal Models, and Current Clinical Practice. Clin Microbiol Rev 31, e00057–17 (2018).

5. Secrieru, A., Costa, I. C. C., O’Neill, P. M. & Cristiano, M. L. S. Anti-malarial Agents as Therapeutic Tools Against Toxoplasmosis—A Short Bridge between Two Distant Illnesses. Molecules 25, 1574 (2020).

6. Bougdour, A. et al. Drug inhibition of HDAC3 and epigenetic control of differentiation in Apicomplexa parasites. Journal of Experimental Medicine 206, 953–966 (2009).

7. Benmerzouga, I. et al. Guanabenz Repurposed as an Antiparasitic with Activity against Acute and Latent Toxoplasmosis. Antimicrob Agents Chemother 59, 6939–6945 (2015).

8. Murata, Y., Sugi, T., Weiss, L. M. & Kato, K. Identification of compounds that suppress Toxoplasma gondii tachyzoites and bradyzoites. PLoS ONE 12, e0178203 (2017).

9. da Silva, M., Teixeira, C., Gomes, P. & Borges, M. Promising Drug Targets and Compounds with Anti-Toxoplasma gondii Activity. Microorganisms 9, 1960 (2021).

10. Kafsack, B. F. C. et al. A transcriptional switch underlies commitment to sexual development in human malaria parasites. Nature 507, 248–252 (2014).

11. Josling, G. A. et al. Dissecting the role of PfAP2-G in malaria gametocytogenesis. Nat Commun 11, 1503 (2020).

12. Waldman, B. S. et al. Identification of a Master Regulator of Differentiation in Toxoplasma. Cell 180, 359-372.e16 (2020).

13. Yasuhara, N., Oka, M. & Yoneda, Y. The role of the nuclear transport system in cell differentiation. Seminars in Cell & Developmental Biology 20, 590–599 (2009).

14. Freitas, N. & Cunha, C. Mechanisms and Signals for the Nuclear Import of Proteins. CG 10, 550–557 (2009).

15. Chahine, M. N. & Pierce, G. N. Therapeutic Targeting of Nuclear Protein Import in Pathological Cell Conditions. Pharmacol Rev 61, 358–372 (2009).

16. Kosyna, F. & Depping, R. Controlling the Gatekeeper: Therapeutic Targeting of Nuclear Transport. Cells 7, 221 (2018).

17. Tay, M. Y. F. et al. Nuclear localisation of dengue virus (DENV) 1–4 nonstructural protein 5; protection against all 4 DENV serotypes by the inhibitor Ivermectin. Antiviral Research 99, 301–306 (2013).

18. Fraser, J. E., Rawlinson, S. M., Wang, C., Jans, D. A. & Wagstaff, K. M. Investigating Dengue Virus Nonstructural Protein 5 (NS5) Nuclear Import. in Dengue (eds. Padmanabhan, R. & Vasudevan, S. G.) vol. 1138 301–328 (Springer New York, New York, NY, 2014).

19. Jans, D. A., Martin, A. J. & Wagstaff, K. M. Inhibitors of nuclear transport. Current Opinion in Cell Biology 58, 50–60 (2019).

20. Walunj, S. B. et al. High-Throughput Screening to Identify Inhibitors of Plasmodium falciparum Importin alpha; Cells 2022, Vol. 11, Page 1201 11, 1201 (2022).

21. Walunj, S. B., Wang, C., Wagstaff, K. M., Patankar, S. & Jans, D. A. Conservation of Importin α Function in Apicomplexans: Ivermectin and GW5074 Target Plasmodium falciparum Importin α and Inhibit Parasite Growth in Culture. IJMS 23, 13899 (2022).

22. Wagstaff, K. M., Sivakumaran, H., Heaton, S. M., Harrich, D. & Jans, D. A. Ivermectin is a specific inhibitor of importin α/β-mediated nuclear import able to inhibit replication of HIV-1 and dengue virus. Biochemical Journal 443, 851–856 (2012).

23. Wagstaff, K. M. & Jans, D. A. Intramolecular masking of nuclear localisation signals: Analysis of importin binding using a novel AlphaScreen-based method. Analytical Biochemistry 348, 49–56 (2006).

24. Dey, V. & Patankar, S. Molecular basis for the lack of auto-inhibition of Plasmodium falciparum importin α. Biochemical and Biophysical Research Communications 503, 1792–1797 (2018).

25. Bhambid, M., Dey, V., Walunj, S. & Patankar, S. Toxoplasma gondii Importin α Shows Weak Auto-Inhibition. Protein J 42, 327–342 (2023).

26. Yammine, A., Gao, J. & Kwan, A. Tryptophan Fluorescence Quenching Assays for Measuring Protein-ligand Binding Affinities: Principles and a Practical Guide. BIO-PROTOCOL 9, (2019).

27. Saini, E. et al. Plasmodium falciparum PhIL1-associated complex plays an essential role in merozoite reorientation and invasion of host erythrocytes. PLoS Pathog 17, e1009750 (2021).

28. Kumar, M., Srinivas, V. & Patankar, S. Upstream AUGs and upstream ORFs can regulate the downstream ORF in Plasmodium falciparum. Malar J 14, 512 (2015).

29. Seeber, F. & Boothroyd, J. C. Escherichia coli β-galactosidase as an in vitro and in vivo reporter enzyme and stable transfection marker in the intracellular protozoan parasite Toxoplasma gondii. Gene 169, 39–45 (1996).

30. Trager, W. & Jensen, J. B. Human Malaria Parasites in Continuous Culture. Science 193, 673–675 (1976).

31. Crabb, B. S. et al. Transfection of the Human Malaria Parasite Plasmodium falciparum. In Parasite Genomics Protocols vol. 270 263–276 (Humana Press, New Jersey, 2004).

32. Knuepfer, E., Napiorkowska, M., Van Ooij, C. & Holder, A. A. Generating conditional gene knockouts in Plasmodium – a toolkit to produce stable DiCre recombinase-expressing parasite lines using CRISPR/Cas9. Sci Rep 7, 3881 (2017).

33. Striepen, B. & Soldati, D. Genetic Manipulation of Toxoplasma gondii. in Toxoplasma Gondii 391–418 (Elsevier, 2007). doi:10.1016/B978-012369542-0/50017-9.

34. Riss, T. L. et al. Cell Viability Assays. in Assay Guidance Manual (eds. Markossian, S. et al.) (Eli Lilly & Company and the National Center for Advancing Translational Sciences, Bethesda (MD), 2004).

35. Bolte, S. & Cordelières, F. P. A guided tour into subcellular colocalisation analysis in light microscopy. Journal of Microscopy 224, 213–232 (2006).

36. Stirling, D. R. et al. CellProfiler 4: improvements in speed, utility and usability. BMC Bioinformatics 22, 433 (2021).

37. Lucantoni, L. et al. A simple and predictive phenotypic High Content Imaging assay for Plasmodium falciparum mature gametocytes to identify malaria transmission blocking compounds. Sci Rep 5, 16414 (2015).

38. Duffy, S., Loganathan, S., Holleran, J. P. & Avery, V. M. Large-scale production of Plasmodium falciparum gametocytes for malaria drug discovery. Nat Protoc 11, 976–992 (2016).

39. Tripathi, A. K., Mlambo, G., Kanatani, S., Sinnis, P. & Dimopoulos, G. Plasmodium falciparum Gametocyte Culture and Mosquito Infection Through Artificial Membrane Feeding. JoVE 61426 (2020) doi:10.3791/61426.

40. Mayoral, J., Di Cristina, M., Carruthers, V. B. & Weiss, L. M. Toxoplasma gondii: Bradyzoite Differentiation In Vitro and In Vivo. in Toxoplasma gondii (ed. Tonkin, C. J.) vol. 2071 269–282 (Springer US, New York, NY, 2020).

41. Lunde, A. & Glover, J. C. A versatile toolbox for semi-automatic cell-by-cell object-based colocalisation analysis. Sci Rep 10, 19027 (2020).

42. Juliana, C. et al. Anti-inflammatory Compounds Parthenolide and Bay 11-7082 Are Direct Inhibitors of the Inflammasome*. Journal of Biological Chemistry 285, 9792–9802 (2010).

43. Sindrewicz, P. et al. Intrinsic tryptophan fluorescence spectroscopy reliably determines galectin-ligand interactions. Sci Rep 9, 11851 (2019).

44. Chan, S. et al. Regulation of PfEMP1–VAR2CSA translation by a Plasmodium translation-enhancing factor. Nat Microbiol 2, 17068 (2017).

45. Yang, S. N. Y. et al. Novel Flavivirus Antiviral That Targets the Host Nuclear Transport Importin α/β1 Heterodimer. Cells 8, 281 (2019).

46. Kosugi, S. et al. Design of Peptide Inhibitors for the Importin α/β Nuclear Import Pathway by Activity-Based Profiling. Chemistry & Biology 15, 940–949 (2008).

47. Kosugi, S. et al. Six Classes of Nuclear Localisation Signals Specific to Different Binding Grooves of Importin α. Journal of Biological Chemistry 284, 478–485 (2009).

48. Stepniewska, K. et al. Efficacy of Single-Dose Primaquine With Artemisinin Combination Therapy on Plasmodium falciparum Gametocytes and Transmission: An Individual Patient Meta-Analysis. The Journal of Infectious Diseases 225, 1215–1226 (2022).

49. Natarajan, K., Singh, S., Burke, T. R., Grunberger, D. & Aggarwal, B. B. Caffeic acid phenethyl ester is a potent and specific inhibitor of activation of nuclear transcription factor NF-kappa B. Proceedings of the National Academy of Sciences 93, 9090–9095 (1996).

50. Takakura, K. et al. Inhibition of nuclear factor-κB p65 phosphorylation by 3,4-dihydroxybenzalacetone and caffeic acid phenethyl ester. Journal of Pharmacological Sciences 137, 248–255 (2018).

51. Coles, V. E. et al. Exploration of BAY 11-7082 as a Potential Antibiotic. ACS Infect. Dis. 8, 170–182 (2022).

52. Abdelhamid Elgendy, W. M., Haggag, Y. A., El-Nouby, K. A., El-Kowrany, S. I. & El Marhoumy, S. M. Evaluation of the effect of guanabenz-loaded nanoparticles on chronic toxoplasmosis in mice. Experimental Parasitology 246, 108460 (2023).

53. D’Alessandro, S. et al. A Plasmodium falciparum screening assay for anti-gametocyte drugs based on parasite lactate dehydrogenase detection. Journal of Antimicrobial Chemotherapy 68, 2048–2058 (2013).

